# Temporal interference stimulation disrupts spike timing in the primate brain

**DOI:** 10.1101/2023.09.25.559340

**Authors:** Pedro G. Vieira, Matthew R. Krause, Christopher C. Pack

**Affiliations:** Montreal Neurological Institute, McGill University Montreal, Quebec, Canada

## Abstract

Electrical stimulation can regulate brain activity, producing clear clinical benefits, but focal and effective neuromodulation often requires surgically implanted electrodes. Recent studies argue that temporal interference (TI) stimulation may provide similar outcomes non-invasively. During TI, scalp electrodes generate multiple electrical fields in the brain, modulating neural activity only where they overlap. Despite considerable enthusiasm for this approach, little empirical evidence demonstrates its effectiveness, especially under conditions suitable for human use. Here, using single-neuron recordings in non-human primates, we show that TI reliably alters the timing of spiking activity. However, we find that the strategies which improve the focality of TI — high frequencies, multiple electrodes, and amplitude-modulated waveforms — also limit its effectiveness. Combined, they make TI 80% weaker than other forms of non-invasive brain stimulation. Although this is too weak to cause widespread neuronal entrainment, it may be ideally suited for disrupting pathological synchronization, a hallmark of many neurological disorders.

## Introduction

Brain stimulation interrogates the relationship between neural activity and behavior, making it an instrumental part of neuroscience research, as well as a tool for treating neurological diseases (Perlmutter & Mink, 2006). However, precise control of neural activity traditionally requires invasive approaches, using electrodes surgically implanted within the targeted brain structure. Since this is both risky and expensive, there has been enormous interest in gaining similar neuroscientific and therapeutic advantages non-invasively. Of particular interest is transcranial alternating current stimulation (tACS) which delivers oscillating electrical current non-invasively, through electrodes attached to the scalp (Figure 1A, yellow line). These currents create electric fields that directly alter single-neuron spike timing, even in the basal ganglia and hippocampus (Krause et al., 2019b; Vieira et al., 2020), with little risk or discomfort for the user. However, because the current is applied to the scalp, targeting deep structures like these with tACS necessarily implies co-stimulation—and indeed, stronger stimulation—of many other brain regions. This has been argued to complicate the interpretation of tACS effects and to limit its translational value (Khatoun et al., 2019; Krause et al., 2019a).

**Figure 1.**
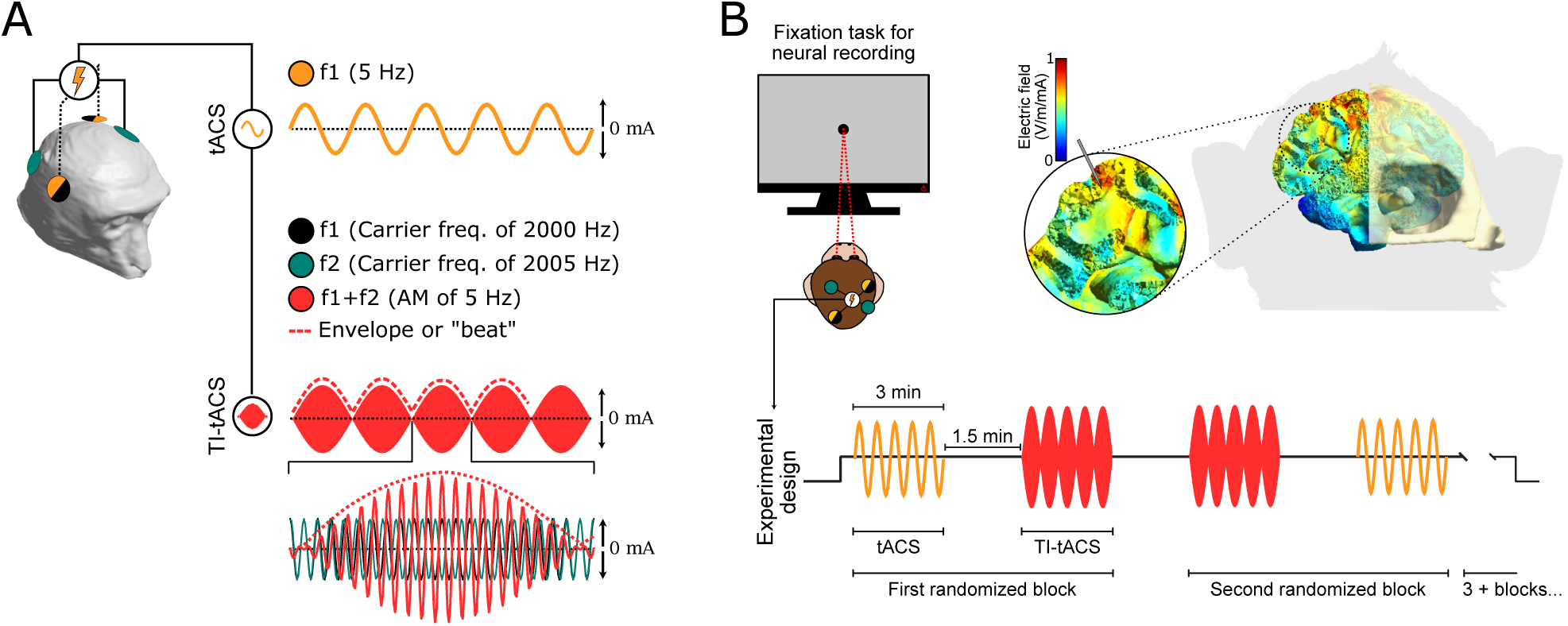
Experimental Overview. A) Schematic depiction of stimulation conditions. During conventional tACS (yellow), the stimulation waveform consists of a sine wave at the target frequency, delivered through a single pair of electrodes. TI-tACS administers high frequency current through two pairs of electrodes, each oscillating at a slightly different frequency (black: 2000 Hz; green: 2005 Hz). Their overlap creates low-frequency AM (red: 5 Hz) where the fields interfere. This component (dashed line) must be extracted through a demodulation mechanism. B) Animals performed a simple visual fixation task while tACS and TI-tACS were administered and single-unit activity was recorded. Electric fields near the recording site were predicted to be no greater than ∼0.7 V/m/mA.

Temporal interference tACS (TI-tACS) attempts to sidestep this problem by exploiting the properties of overlapping electric fields (Grossman et al., 2017). Instead of delivering alternating current at the intended frequency, TI-tACS creates two electric fields, each using a separate pair of electrodes and each oscillating at a slightly different carrier frequency (Figure 1A, black and green lines). Individually, the carriers oscillate too rapidly to affect neural activity. However, wherever the two electric fields overlap (Figure 1A, red), their interference produce an amplitude modulation (AM); this phenomenon is sometimes called a “beat” or “interference pattern” as well. The AM fluctuates at the difference between the carrier frequencies: for example, superimposing 2000 Hz and 2005 Hz carriers creates a 5 Hz AM. Judicious configuration of the electrodes could therefore produce AM that is largely confined to the target brain structure and oscillating in a physiologically relevant frequency range (<100 Hz).

It is tempting to imagine that the resulting AM acts just like conventional tACS—and it is often claimed that it does; see Mizakhalili (2020) for a discussion of this issue — but the underlying mechanism must be fundamentally different. The AM is the outline, or envelope (Figure 1A, red dashed line), of the signal produced by the two overlapping carriers and must be extracted from them. This procedure is called demodulation and it is unclear how efficiently neurons can perform it (Mirzakhalili et al., 2020), how they might do so (Luff et al., 2023; Rosenberg & Issa, 2011), and there is some question as to whether they do it at all (Iszak et al., 2023). Existing neurophysiological data demonstrating the effectiveness of AM come from experiments using electric fields a thousand times stronger (Rampersad et al., 2019) than those which are safe for human use (Cassarà et al., 2022). Moreover, it can be shown mathematically that demodulation requires non-linear operations, making it challenging to interpret existing neurophysiological data. This has led some to question whether TI-tACS is feasible in humans at all (Iszak et al., 2023).

Here, we measure the effects of TI-tACS on neurons in alert non-human primates, which are anatomically and physiologically similar to humans. Under stimulation conditions that closely match human use, we find that TI-tACS predominantly affects spike timing and not spike rate, just as has been observed with conventional tACS (Krause et al., 2019b, 2022). However, the effects of TI-tACS appear to be substantially weaker than those of tACS, due to greater shunting of current away from the brain and incomplete demodulation of the AM. As a result, TI-tACS largely disrupts endogenous rhythms but fails to impose new ones, suggesting that its primary therapeutic value will be to reduce synchronous oscillations in deep brain structures.

## Results

We recorded the activity of 234 neurons in two non-human primates. During these recordings, the animals performed a simple visual fixation task to control extraneous sensory and cognitive factors that might otherwise affect neural activity (Fig 1B; Methods: Behavioral Task). In each session, we delivered TI-tACS and tACS in a randomized block design, allowing us to compare the effects of different types of brain stimulation against unstimulated baseline activity and against each other. Stimulation was delivered through electrodes placed on the scalp at locations chosen to produce effective TI-tACS stimulation of our recording sites (see *Methods*). For TI-tACS, one carrier frequency was fixed at 2000 Hz, and the other varied across experiments between 2005, 2010, and 2020 Hz, producing AM frequencies of 5 Hz – 20 Hz, covering the range used in most human studies. Field strengths were also similar to human studies: according to our finite-element model (see *Methods)* the upper bound in our experiments was no greater than 0.7 V/m, in the middle of the 0.4 – 1.0 V/m range estimated for humans (Jackson et al., 2016). As detailed in the Methods, we performed a number of technical controls to verify that our equipment generated the intended stimuli and did not produce artifacts affecting measures of neural activity (Figure S1).

### TI-tACS affects spike timing, not rates

In principle, TI-tACS could affect neural activity in two ways. First, it might change the timing of spikes relative to ongoing oscillations, as observed previously for tACS (Krause et al., 2019b). Alternatively, the overall rate of spiking activity could change, indicating an effect on excitability (Chaieb et al., 2011) or a conduction block; the latter has been proposed as a mechanism by which TI-tACS could decrease firing rates (Kilgore & Bhadra, 2014).

Figure 2 shows the activity of two example neurons under baseline conditions (blue) and during the application of TI-tACS (red). The neuron in Figure 2A fired more rhythmically in response to TI-tACS, with action potentials preferentially occurring during the rising phase of the 20 Hz envelope. We quantified this rhythmicity with the phase-locking value (PLV), a quantity that indexes the consistency of firing across an LFP or stimulation cycle (see Methods). A PLV of 0 corresponds to completely unstructured firing, while a PLV of 1 indicates that a neuron is completely entrained by the oscillation and therefore fires at precisely the same phase of each cycle.

**Figure 2.**
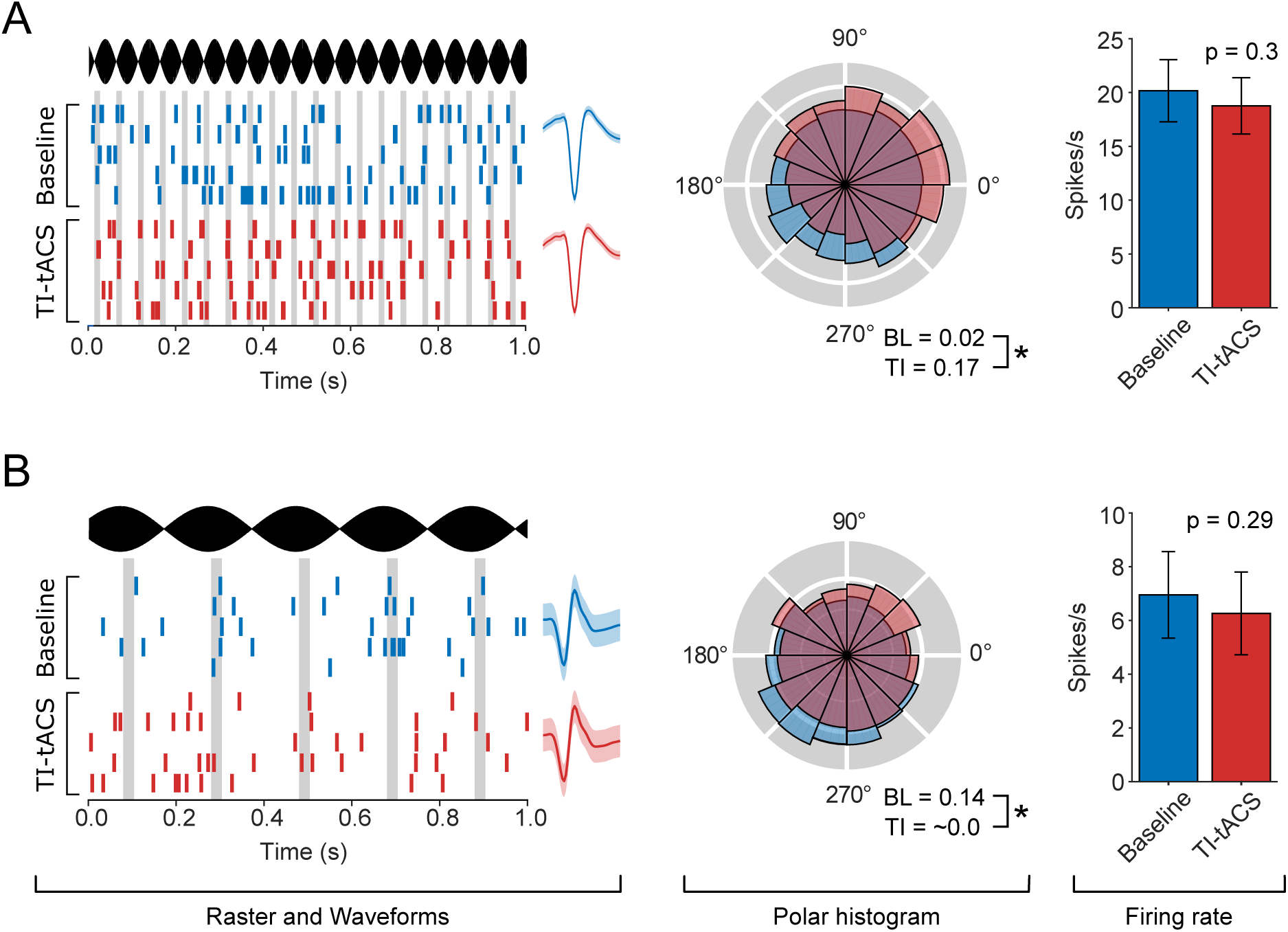
Example neurons receiving TI-tACS. A) Example of a neuron entrained by TI-tACS. The left column contains raster plots from 5 1-s segments of data, showing the AM waveform (black) and the timing of spikes emitted during TI-tACS (red) and baseline (blue). Vertical grey lines indicate the preferred phase during TI-tACS. The average spike waveforms during each condition are shown, to demonstrate that single-unit isolation was maintained. The center column contains spike density histograms showing the relative probability of spiking at each phase of the TI-tACS (red) or baseline LFP (blue), which are summarized by PLV values (See Methods). The right column compares average firing rates across conditions. No significant difference was observed. B) An example neuron desynchronized by TI-tACS, plotted in the same style as Panel A. * indicates p < 0.01

The neuron in Figure 2A showed a significant increase in PLV from 0.02 at baseline to 0.17 during TI-tACS (p < 0.01; randomization test). This increase is evident in the polar histogram (middle panel), which shows a concentration of spikes at phases between 0° and 90° during TI-tACS (red), but little preference for any phase of the baseline LFP oscillation (blue). In contrast, the neuron in Figure 2B became less rhythmic by the application of 5 Hz TI-tACS, as indicated by the significant decrease in PLV from 0.14 during baseline to approximately 0 during TI-tACS (p < 0.01; randomization test). For this neuron, the baseline preference for spiking at LFP phases around 210° (middle panel, blue) was eliminated by the application of TI-tACS (red). For both neurons, there was no detectable difference in the spike rate between stimulation conditions (right panels). As shown by the waveforms (left panels), neurons were well isolated within and across conditions, suggesting that signal loss cannot account for these results (see also *Methods*: *Validation*).

We performed similar measurements under a wide range of conditions, using recordings from multiple cortical areas (V4, 7A, and MT) and stimulation with multiple AM frequencies (5, 10, and 20 Hz) and carrier amplitudes (±1.0, ±2.0, and ±2.5 mA). To summarize these experiments, Figure 3 shows the change in PLV (ΔPLV = Stimulation – Baseline) for each of the 234 neurons in our sample. TI-tACS had clear effects on spike timing, causing individually significant changes in 28% (65/234) of these neurons, shown here in red (p<0.05; per-cell randomization tests). Sixteen neurons, including the example cell in Figure 2A (indicated here by a star), showed individually significant increases in entrainment. However, decreased entrainment was a much more common outcome, found in 75% (49/65) of the TI-tACS responsive neurons, including the example in Figure 2B (diamond).

**Figure 3.**
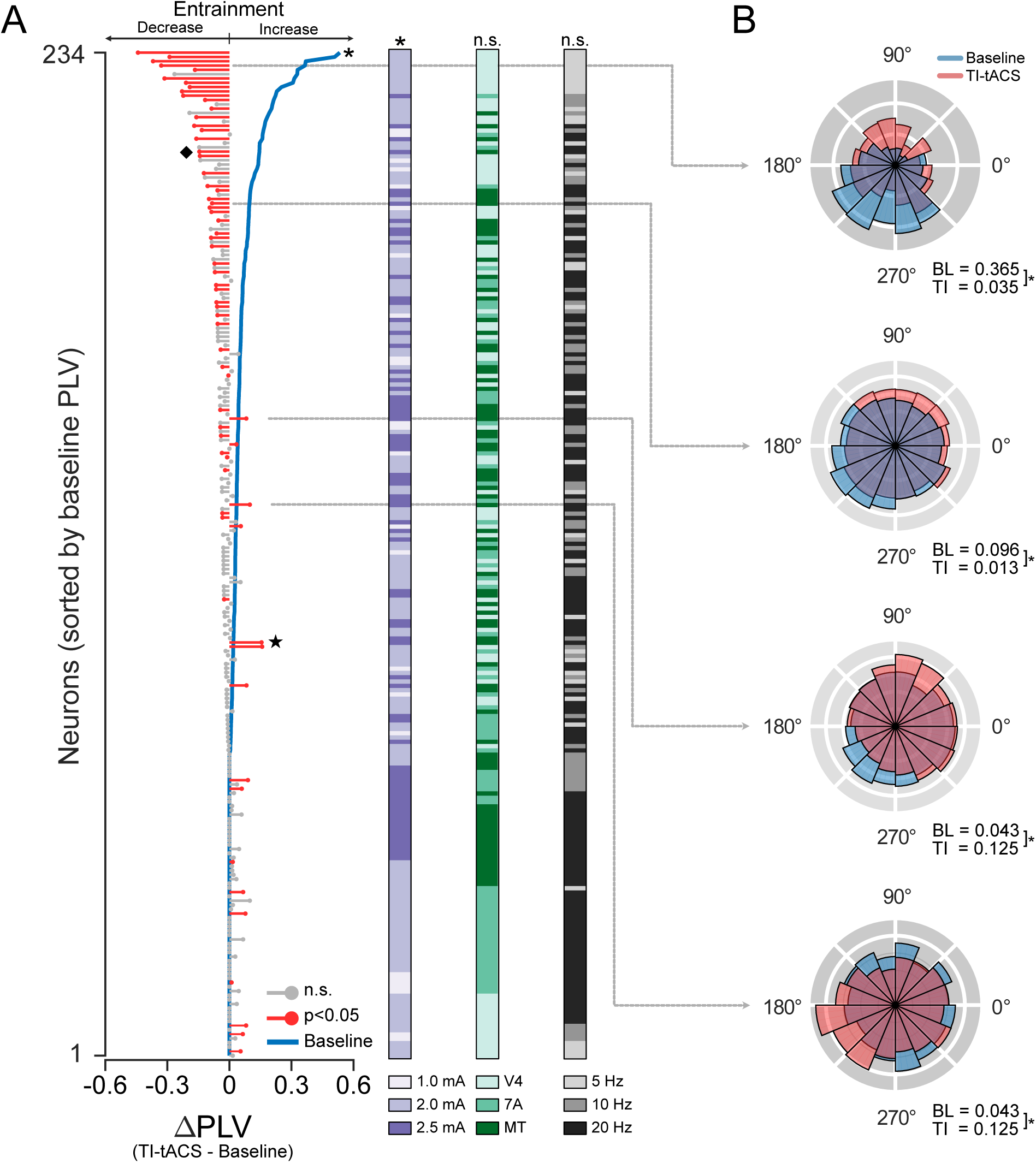
TI-tACS competes with baseline activity. A) Population results during TI-tACS. Each line indicates how a single neuron’s spike timing changed due to TI-tACS (ΔPLV = TI-tACS – Baseline). Red lines indicate individually significant changes (p < 0.05); grey lines were not significantly altered on a per-cell basis. Neurons were sorted by baseline PLV (blue line). Other experimental conditions are indicated by the colored bars (violet: carrier amplitude; green: brain area, grey: AM frequency). * and n.s. (not significant) refer to the results of the ANCOVA described in the text; See also Figure S2 for individual conditions. B) Spike density histograms for four example cells, plotted in the same style as Figure 2, and exhibiting a range of effects. The star indicates the example neuron in Figure 2A; the diamond denotes the cell from Figure 2B.

Prior work with conventional tACS has found similar bidirectional effects (Asamoah et al., 2022; Krause et al., 2022). These effects are thought to reflect competition between the stimulation and ongoing brain activity for control over spike timing: when baseline entrainment is weak, external stimulation completely imposes new rhythms on the spike train. In the presence of stronger baseline entrainment, the same external stimulus cannot dominate spike timing; instead, both vie for control over when the neuron spikes and the overall result is less rhythmic spiking (Krause et al., 2022). To determine whether similar competition occurs with TI-tACS, we sorted the cells in Figure 3 by baseline PLV (blue line). We found a strong negative correlation between baseline PLV and ΔPLV (Spearman’s ⍴= –0.43; p << 0.001) using Oldham’s method to correct for regression to mean. An ANCOVA confirmed that baseline PLV, stimulus amplitude, and their interaction were the only significant predictors of ΔPLV (Baseline PLV: p << 0.001; F(1)=222.58; Amplitude: p=0.033; F(2)=3.46; Interaction: p=0.037: F(2)=3.33). Stimulation frequency and brain area had no effect on ΔPLV, so we pooled data these conditions for the remaining analyses.

Unlike the dramatic changes in spike timing, TI-tACS caused only minor changes in firing rates. The median firing rate during baseline blocks was 4.1 Hz (95% CI: 3.4—5.1 Hz), while the median during TI-tACS was 3.9 Hz (95% CI: 3.3 – 4.5 Hz). An ANCOVA found that firing rates during TI-tACS were predicted by the baseline firing rate (F(1) =1297; p<<0.001) but not the stimulation amplitude, frequency, or brain area (F(2) > 2.2; p>0.12). The coefficient for the baseline amplitude was very nearly 1.0 (95% CI: [0.77-0.99]), suggesting that firing rates were generally unchanged. In an exploratory analysis, we did find small but significant effects in the subpopulation of neurons that received 5 Hz AM, but these were very small (∼0.2 Hz) and on the cusp of significance (p=0.03; Wilcoxon Sign-Rank Test). Overall, sixteen percent of cells showed individually significant differences in firing rates across conditions (p <0.05; mixed-effect model), but the average magnitude of these changes was not significantly different from zero (p=0.85; 1-sample t-test). Thus, any changes in firing rate with TI-tACS were likely to be quite modest and are not consistent with a widespread conduction block or a consistent increase in excitability.

### Conventional tACS is stronger than TI-tACS

These data suggest that TI-tACS and tACS affect neurons similarly. To compare the two modalities more directly, we recorded responses to both forms of stimulation in a subset of 154 neurons. The tACS frequency and TI-tACS AM frequency were matched within each cell: for example, 5 Hz tACS was compared against 2000+2005 Hz TI-tACS, which produces a 5 Hz AM. We held the tACS amplitude constant at ±1 mA, while varying the amplitude of both TI-tACS carriers across recordings.

Figure 4A shows the individual results for the 65 neurons that were significantly modulated by either TI-tACS (red) or conventional tACS (yellow) (randomization tests; p<0.05). The pattern of effects was broadly similar. Both modalities decreased entrainment in the presence of highly structured baseline activity (blue line, top), and both increased entrainment for neurons with baseline PLVs near zero (blue line, bottom). However, at intermediate levels of baseline activity, we sometimes observed conflicting effects. The example neuron in Figure 4B showed increased entrainment for ±1 mA tACS at 10 Hz (yellow), but significantly decreased entrainment by TI-tACS at the same AM frequency (red). This occurred even though the amplitude of the TI-tACS carrier was much larger (±2.5 mA) than the current used for conventional tACS (±1 mA).

**Figure 4.**
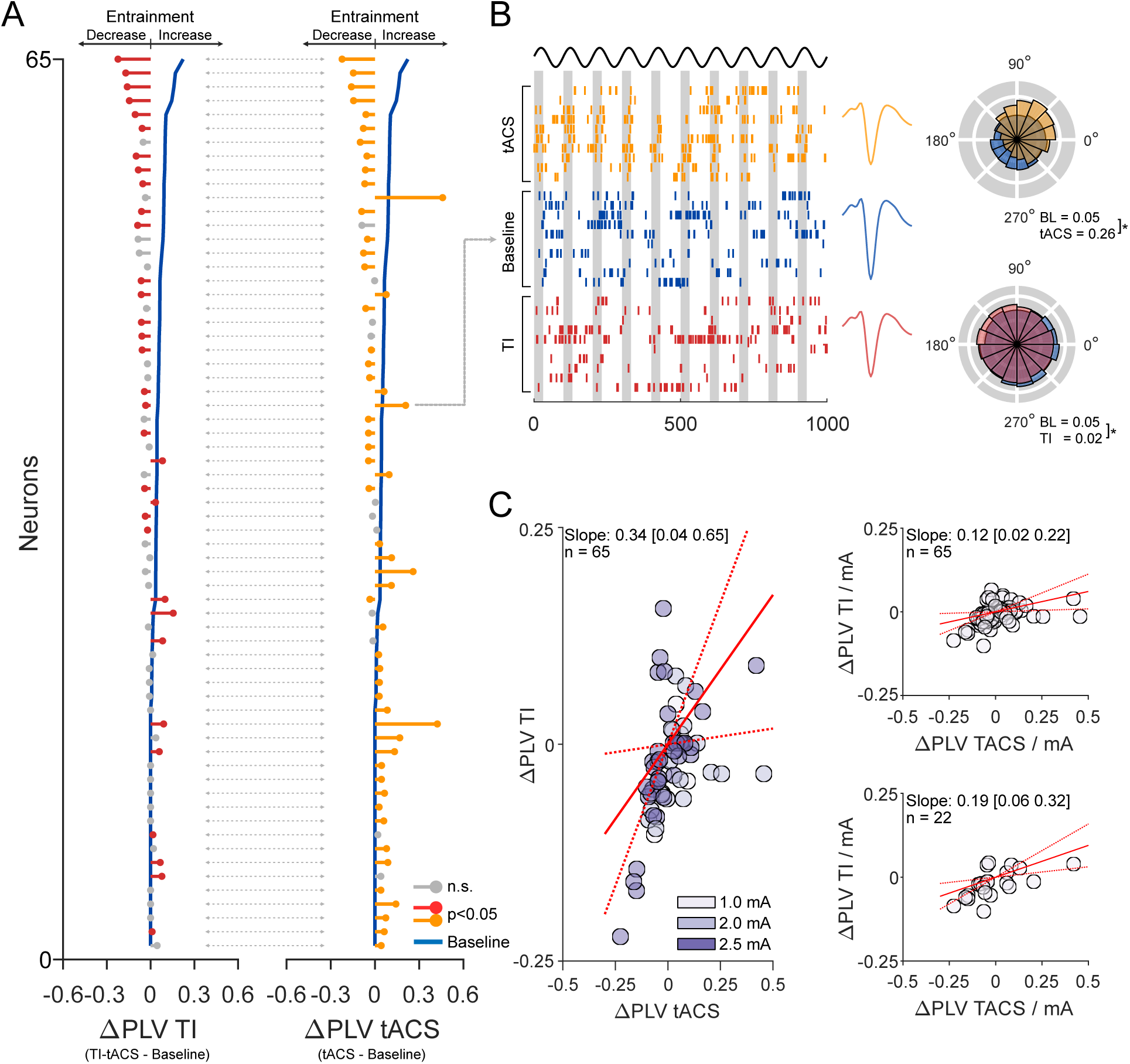
TI-tACS is weaker than conventional tACS. A) Data from 65 neurons where either TI-tACS or tACS had a significant effect on PLV (either direction; p < 0.05), plotted in the style of Figure 3A. B) At intermediate levels of background activity, tACS and TI-tACS sometimes had opposing effects on spike timing. For this cell, plotted in the style of Figure 2, tACS significantly entrained neural activity but TI-tACS significantly desynchronized it (p<0.05; randomization tests). C) PLV changes during TI-tACS plotted against those during TI-tACS. Colors indicate the current amplitude during TI-tACS. The solid red line indicates the best-fit slope; dashed red lines denote the limits of the 95% confidence interval. Considerably more current was used during TI-tACS so we scaled the TI-tACS ΔPLV to account for this and reanalyzed the data (inset, top). The bottom inset plot repeats the analysis using only the 22 neurons which were significantly modulated by both (rather than either) modalities. All three analyses suggest that TI-tACS is weaker than conventional tACS.

These data suggest that conventional tACS may be stronger than TI-tACS, and therefore more capable of imposing a rhythm onto the neuron’s spike train. For the cell in Figure 4B, this would be reflected as an increased PLV as well as a shift in preferred phase towards the first quadrant (0-90°), as suggested by models of the underlying biophysics (Radman et al., 2007). At the population level, this hypothesis suggests that tACS should increase entrainment in more neurons. This was in fact the case, with conventional tACS entraining significantly more neurons (30/65, or 46%: 95% CI: [39%-59%]) than TI-tACS (11/65 neurons, 17%).

Figure 4C formalizes this comparison by comparing the relative effects of conventional tACS and TI-tACS on these 65 neurons, using Deming regression to account for the uncertainty in both ΔPLV measurements. In absolute terms, the slope of the best-fit line suggests that TI-tACS is 0.34 times the strength of conventional stimulation (95% CI: [0.04-0.65]). However, this does not account for the increased current used in some TI-tACS experiments, which was in some cases delivered at ±2.0 or ±2.5 mA per pair rather than the ±1 mA used in all conventional tACS blocks. We therefore scaled the PLV values by the current delivered through a pair of electrodes, under the assumption that, at least over a limited range, stimulation effects are roughly proportional to the amount of current applied (Johnson et al., 2020). On this adjusted per-mA basis, the effects of TI-tACS were on average 12% (Deming regression; 95% CI: 2-22%) of those caused by tACS (Figure 4C, top). Restricting this analysis to cells that were significantly modulated by both (rather than either) condition yielded similar results (Figure 4C, bottom).

### Why is TI-tACS weaker?

A combination of three mechanisms can explain why TI-tACS is weaker than conventional tACS.

First, much of the current in the TI-tACS stimulation never reaches the brain but is instead shunted through skin, muscle, and bone. Although this limits all forms of tES (Vöröslakos et al., 2018), biological tissue exhibits frequency-dependent changes in conductivity (Hasgall et al., 2022) that may specifically affect TI-tACS. We assessed the potential impact of carrier frequency by measuring the size of the intracranial stimulation artifact at frequencies ranging from 10-2000 Hz. Figure 5 shows that the stimulation artifact exponentially decreases with increasing stimulation frequency in the brain (black line). Since this pattern was not observed in control experiments with our recording system (grey line), it likely has a biological origin (*Methods*: *Validation*). Thus, the well-known shunting effects of transcranial stimulation might be particularly problematic for the high-frequency carriers used in TI-tACS.

**Figure 5.**
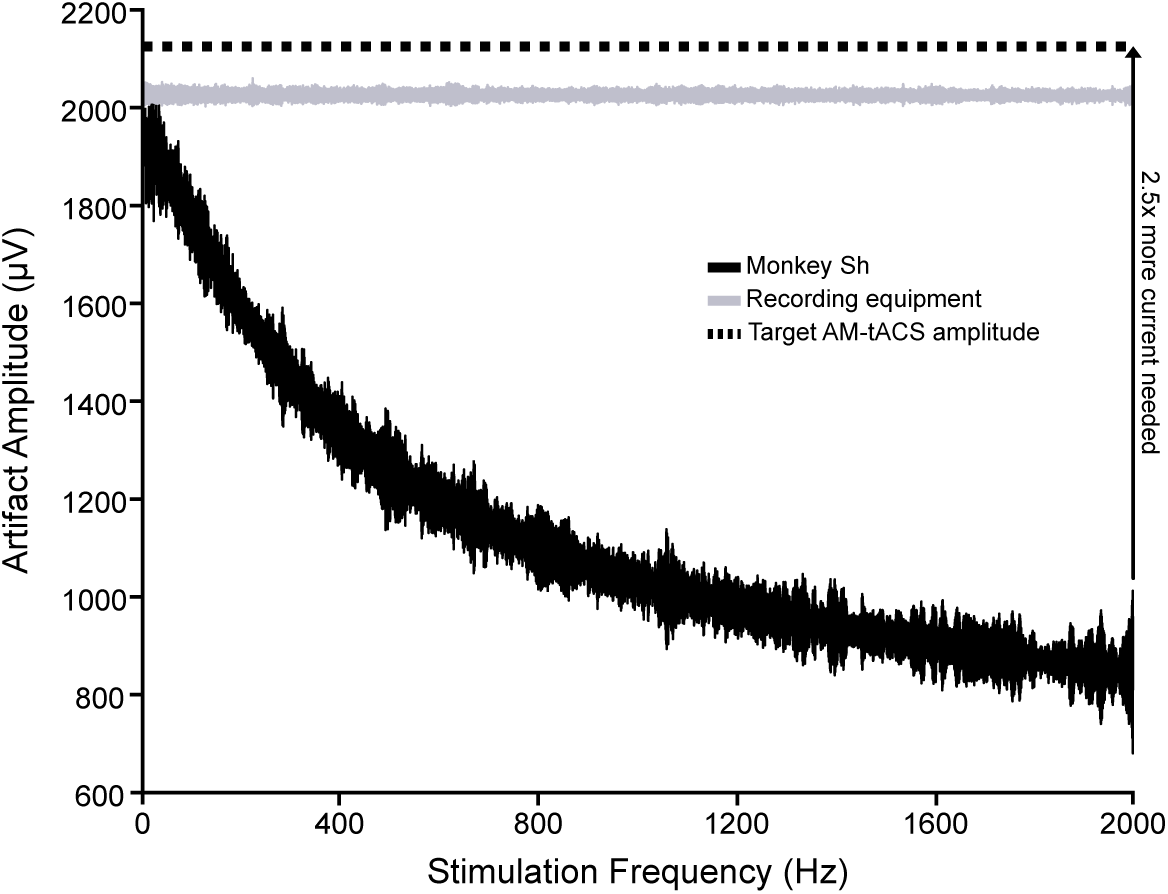
Frequency-dependent conductivity limits TI-tACS. The amplitude of the stimulation artifact was measured at frequencies from 10-2000 Hz. In the brain, this shows a clear decay with increasing frequency (black). This effect is not due to the properties of our recording system, as applying the same stimulus to the head stage shows minimal decay (gray). This curve was also used to calibrate the AM-tACS experiment in Figure 6, so that its 2000 Hz carrier produced the same modulation depth as conventional low-frequency tACS.

Second, neurons may be unable to fully extract the AM component of the TI-tACS stimulation (Mirzakhalili et al., 2020). To test this possibility directly, we performed an additional experiment in which we delivered both conventional tACS and a single amplitude-modulated sine wave (AM-tACS) through the same pair of electrodes (Figure 6A) and at the same frequency (10 Hz vs. 10 Hz AM). We compensated for the loss of current with high-frequency carriers by adjusting the AM-tACS stimulation so that it produced similar (in fact slightly larger) modulation depths as the tACS stimulation (see *Methods* and Figure 5). Consequently, differences in entrainment between the two conditions must be due to imperfect extraction of the AM waveform.

**Figure 6.**
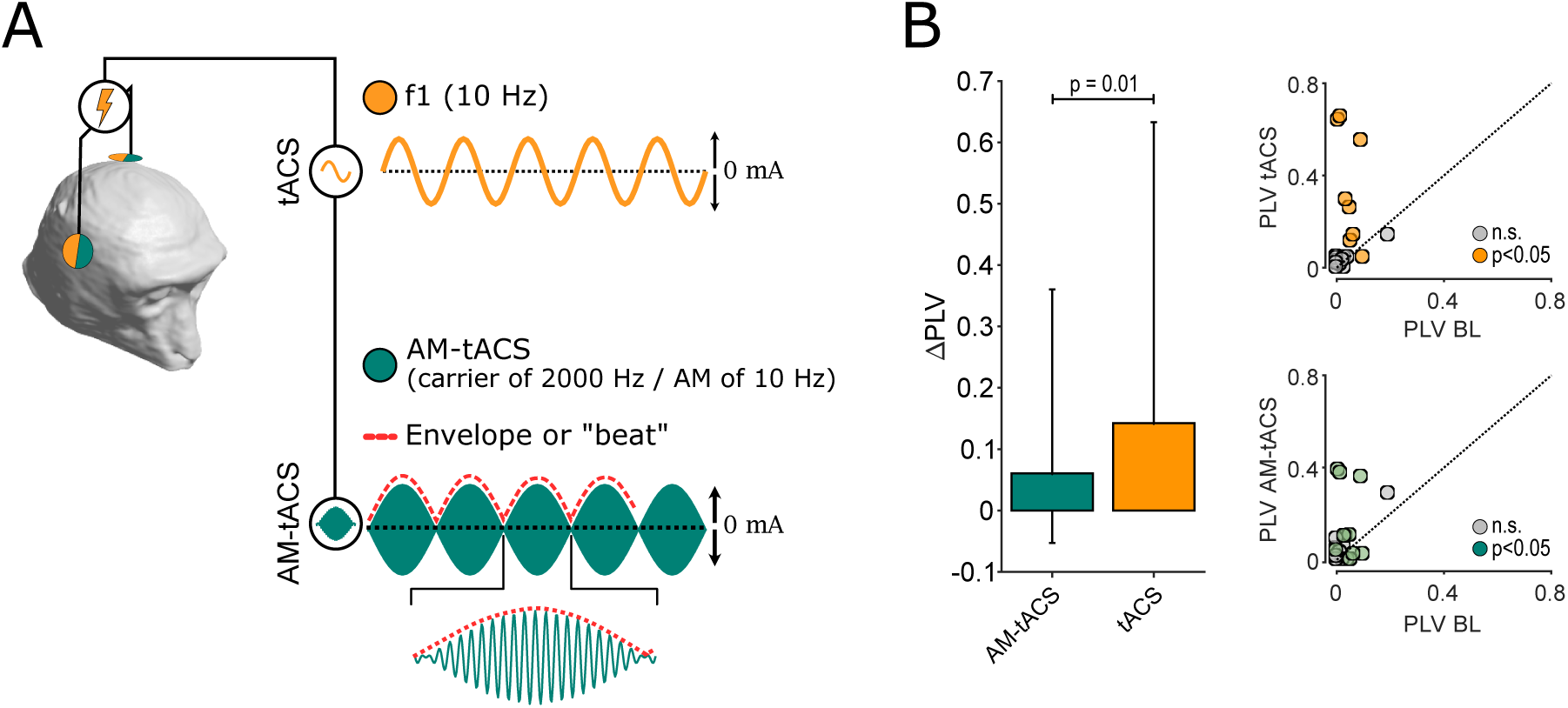
Demodulation losses limit TI-tACS. A) Schematic depiction of AM-tACS experiments, as in Figure 1A. Unlike TI-tACS, both conventional tACS (yellow) and AM-tACS (green) are delivered through the same two electrodes. The AM carrier uses the same frequency as TI-tACS: 2000 Hz with an AM frequency (red) matching the tACS. B) Medians ΔPLVs for responsive neurons during ±2.5 mA AM-tACS (green), and ±1.0 mA tACS (yellow). Scatter plots show the effects vs. baseline (BL) for each condition.

Stimulation with either modality had a significant effect on the entrainment of half the cells (10/20; 50%), seven of which were significantly modulated by both (Figure 6B). Since non-response could reflect biophysical factors like orientation relative to the electric field (Radman et al., 2009) or limited statistical power, we focused on the latter group of responsive cells. Against a low level of baseline entrainment (median PLV = 0.043), conventional tACS increased the median PLV to 0.20 (95% CI: [0.043 – 0.64]). By comparison, AM-tACS only increased the median PLV to 0.10 (95% CI: [0.023 – 0.38]). This two-fold difference between the two conditions was significant (p < 0.009; Wilcoxon sign-rank test), suggesting that neurons are poor demodulators of AM signals.

Finally, due to the underlying physics, the AM depth produced by two interfering electric fields is capped by the magnitude of the weaker one (see Appendix B and E of Huang & Parra, 2019). The greatest focality is usually achieved by increasing the separation of the electrode pairs on the scalp. However, this also requires moving the electrodes away from the optimal scalp location, which weakens the field strength. These problems may be somewhat greater in our animal models, where the heads are smaller, and some potential scalp locations are blocked by the experimental apparatus. Nevertheless, it is expected to occur for many targets in humans as well (Huang & Parra, 2019).

## Discussion

Here, we have shown that TI-tACS non-invasively alters spike timing in primates (Figures 2-3). This suggests that neurons are capable of demodulating amplitude-modulated electric fields, which can in turn increase or decrease the prevalence of rhythmic spiking. However, this demodulation is inefficient (∼50%; see Figure 6). The effects of TI-tACS are further limited by the high frequency of the carrier (Figure 5) and the need for multi-electrode stimulation to create the interference pattern. Combined, these factors make TI-tACS approximately 80% weaker than conventional tACS using the same amount of current (Figure 4).

Consequently, TI-tACS in humans is unlikely to be strong enough to completely overcome most ongoing activity and impose new rhythms onto it (Krause et al., 2022). Instead, it will tend to promote more uniform, less rhythmic spiking through competition with ongoing brain activity. We emphasize that this is not a categorically different effect from conventional tACS, which exhibits similarly graded competition. The stimulation is simply weaker during TI-tACS.

Our work was purposely conducted with currents that generate intracranial electric fields comparable to those used in previous human studies (Figure 1B). This is in contrast to prior animal work, which used much stronger stimulation that is unlikely to be tolerable or safe in humans. Grossman et al. (2017) applied ±0.5 mA TI-tACS to mice and found that it elicited spiking activity, as revealed by widespread cFos expression near the stimulation focus. However, this current was delivered to a small animal and through a thinned skull, and therefore produced electric fields of between 60 and 383 V/m (Grossman et al., 2017; Rampersad et al., 2019), orders of magnitude stronger than the 0.4 – 1.0 V/m typical in humans (Jackson et al., 2016). Older work by Kuzin et al. (1965) was done in humans but used current sufficient to induce immediate loss of consciousness. In these experiments, the powerful stimulation overcame any inefficiencies in the transfer of current or its demodulation. However, they do not accurately reflect how TI-tACS acts during human neuromodulatory use, where the applied currents are much weaker. As with the past confusion about the effects of conventional tACS (Liu et al., 2018), these results highlight the importance of using realistic conditions to probe the mechanisms of human brain stimulation. Indeed, our results do seem consistent with recent experiments in humans which found that TI-tACS targeted at the hippocampus decreased BOLD activity (Violante et al., 2022).

It might be possible to improve the effectiveness of TI-tACS by moderately increasing the strength of TI-tACS induced electric fields. Most existing devices limit currents to a maximum of ±2.5 mA, based on safety and tolerability considerations derived from conventional tACS. Both safety and tolerability increase at higher stimulation frequencies, so up to 7 mA of stimulating current might be acceptable during TI-tACS (Cassarà et al., 2022). The fields created by this stronger stimulation would be approximately threefold stronger, which would likely be sufficient to reliably increase single-neuron entrainment, but still insufficient to alter spike rate (Krause et al., 2022; Johnson et al., 2020).

Alternatively, more effective TI-tACS could be achieved with TI-tACS waveforms that are more readily demodulated by the brain. The mechanisms responsible for demodulation are not yet well-understood, but they likely involve a combination of cell-intrinsic and network properties. Within a single cell, asymmetries between inward-going sodium and outward-going potassium channels may be sufficient to generate susceptibility to AM electric fields (Mirzakhalili et al., 2020; Plovie et al., 2023). Within larger networks, frequency-dependent adaptation of a circuit could also contribute to demodulation (Esmaeilpour et al., 2021). Sensory systems selective for AM also use mechanisms across these scales, with AM demodulation occurring within a single cell in electrosensory systems (Metzen & Chacron, 2015), across distributed retinocortical networks in the visual system (Baker & Mareschal, 2001; Rosenberg & Issa, 2011), and via a combination of both in the auditory system (Shofner et al., 1996). These mechanisms share common nonlinear operations like rectification, which are also used in engineered systems like radios. However, they have different biophysical implications, and as such this could be a fruitful topic of further research.

A biophysical consideration apparent in our data is that the strength of transcranial stimulation depends on the stimulation frequency. In most experiments, especially in humans, this cannot be measured directly but is instead estimated from finite-element models of the participants’ heads, like the one used to position our electrodes (see *Methods).* These incorporate conductivity estimates for each element of the head (e.g., bone or grey matter), but the estimates are often taken from sources using low frequencies or direct current. While this approximation seems valid for conventional tACS (Huang et al., 2017), the substantial frequency-dependent changes we observed influence the strength of electric fields reaching the brain during TI-tACS (Figure 5). Future experiments should therefore consider the specific carrier frequencies, as conductivity values vary substantially even between 1 and 2 kHz (Hasgall et al., 2022). Corrected models and field strength measurements in human subjects, when combined with our data, may offer a more realistic portrait of TI-tACS’s effects in humans.

Our data suggest that there is little rationale for using AM-tACS, which was initially proposed to have two main advantages over conventional tACS: facilitating concurrent EEG by minimizing spectral overlap between the stimulus waveform and targeted brain oscillation and reducing user discomfort. In practice, however, AM-tACS may still cause signal processing confounds, especially in lower-bandwidth equipment used for EEG (Kasten et al., 2018 but note this was not present in our apparatus; see Figure S1). Human participants do report that AM-tACS is more tolerable, in terms of pain sensations and phosphene production than conventional tACS, but these effects are still not completely eliminated (Thiele et al., 2021). However, AM-tACS’s effects on spike timing were only half as strong as conventional tACS (Figure 6), consistent with prior in vitro work (Esmaeilpour et al., 2021). Consequently, this technique seems to have all the drawbacks and none of the advantages of other forms of tACS.

Recent work has also proposed adapting temporal interference to other modalities, especially transcranial magnetic stimulation (TMS) (e.g., Khalifa et al., 2023; Labruna et al., 2023). The neurophysiological effects of these methods have not yet been fully characterized, but users will still need to contend with similar issues related to sub-optimal electrode/coil placement and incomplete demodulation. On the other hand, TMS produces stronger electromagnetic fields, which may overcome some of these inefficiencies.

These limitations aside, TI-tACS, as delivered through existing technologies, could prove useful in certain settings. As we have shown here, the weak electrical stimulation delivered by TI-tACS tends to desynchronize neurons, and a non-invasive method for producing targeted desynchronization may be therapeutically valuable. Excess synchrony has been implicated in a wide range of pathological conditions, including epilepsy, Parkinson’s Disease (McConnell et al., 2012), and schizophrenia (Venables et al., 2009). Even in healthy brains, desynchronized states are often associated with improved information processing (e.g., selective attention; Cohen & Maunsell, 2009), because they allow neurons to act independently, improving the richness of the neural code. This kind of desynchronization is thought to be the mechanism of action for many existing treatments, including DBS (McConnell et al., 2012). Our results suggest that TI-tACS may be able to desynchronize neural activity simply and non-invasively.

## Acknowledgments

We thank Dr. Fernando Chaurand, Julie Coursol, Cathy Hunt, and Jessica Hutta for outstanding technical assistance and Tudor Sintu for help with the finite-element modelling. Thank you to Klaus Schellhorn (neuroConn GmbH, Ilmenau) and Dr. Stephen Frey (Rogue Research Inc., Montreal) for providing the TI-tACS equipment. This work was supported by grants from NSERC (DH-2022-00476) and the CIHR (202104PJT-461642).

## Materials and Methods

### Experimental Approach

We collected data from two adult macaque monkeys: Monkey Sa (an 11 year old male; 14 kg) and Monkey Sh (10 year old female; 8.4 kg). Apart from the temporal interference stimulation, the approaches and techniques are very similar to those used in our previous work but are nevertheless described in detail below. Monkey Sa also participated in some of those experiments, but the data presented here has not been reported previously; Monkey Sh was obtained specifically for these experiments.

### Ethics Statement

The Montreal Neurological Institute’s Animal Care Committee approved these experiments under Animal Use Protocols #1870 and #5031. The work was also supervised by the Institute’s veterinarian and trained animal health staff and followed additional guidelines from the Canadian Council on Animal Care and the Weatherall Report on the Use of Non-Human Primates in Research. Specifically, the animals received varied and daily environmental enrichment, both in their home cages and through regular access to a large play arena. Since animals were socially housed, they also had visual and tactile access to conspecifics when not in the lab.

### Animal Preparation

We first obtained high resolution T1– and T2-weighted MRIs of each animal’s head and neck, which were used for surgical planning and to optimize the tES montages (see below). T1w images were acquired with an MP-RAGE sequence (TR = 2300 ms and TE = 3.59 ms); the T2 images used a TR = 2800 ms and TE = 489 ms instead. Between 7 and 10 separate volumes consisting of 0.57mm isotropic voxels were acquired for each pulse sequence. Volumes were denoised, aligned, and averaged with FSL and AFNI. Titanium head holders (Hybex Innovations, Montreal) were then attached to each animal’s skull using standard surgical techniques. After an eight-week recovery period, animals were familiarized with the laboratory, head restraint, and the behavioral task.

To prepare animals for neurophysiological recording, we then attached a set of MR-opaque fiducial markers to the animals’ headposts and acquired a second set of T1w MRIs. After manually masking regions obscured by susceptibility artifacts due to the headposts, the two scans were aligned with each other and the NMT Atlas (Seidlitz et al., 2018). We then performed a second surgery to implant a recording chamber (Crist Instruments, Hagerstown, Maryland, USA). The chamber was implanted at an oblique angle, so as to provide access to multiple visual areas, including 7A, MT, and V4v, using a frameless sterotaxic neuronavigation system (Brainsight Vet, Rogue Research). Bone within the chamber was carefully removed, leaving the underlying dura intact. In monkey Sa, the trajectory was verified with an additional postoperative MRI. For monkey Sh, we instead confirmed the trajectory via the visual response properties of neurons in each area and the white/grey matter transitions visible on both the MRIs and neurophysiological recordings.

### Behavioral Task

Changes in arousal, motivational state, and oculomotor activity can strongly affect rhythmic brain activity—and thus the effects of tES as well. We therefore used a simple visual fixation task to ensure that animals remained in a consistent behavioral state during the experiment. The animals sat in an enclosed primate chair (Crist Instruments, of Hagerstown, Maryland) in a dark, copper-lined testing booth. A computer monitor was placed 57 cm in front of them, covering the central 30 ° x 60 ° of their visual field. Using an infrared eye tracker (EyeLink II; SR Research) to monitor gaze position, animals were operantly trained to fixate a small dark target (0.5° radius), presented against a neutral grey background (54 cd/m^2^). Fruit juice rewards were given whenever their eyes remained within 1-2° of the fixation target for 1–2 seconds. The exact interval between successive rewards was drawn from an exponential distribution to prevent possible entrainment to rewards and expected rewards. Testing sessions typically lasted 60-120 minutes but were discontinued early when animals consistently failed to maintain fixation. Custom software, written in Matlab (The Mathworks, Natick, MA) controlled the behavioral task and coordinated the eyetracker, stimulator, and recording equipment.

### Neurophysiology

Daily-acute recordings were made through the recording chambers implanted on the skull. After penetrating the dura with a 22-gauge stainless steel guide tube, we inserted 32 channel linear arrays (V-Probe; Plexon, Dallas, Texas, USA). Each site on the array had an impedance of 200—400 kΩ; adjacent sites were separated by 150 µm. Arrays were then lowered into the target structures by a computer-controlled microdrive (NAN Instruments, Nazareth Illit, Israel). Targets were identified using depths derived from the MRI and neuronavigation system as well as patterns of activity.

Signals were recorded from these electrodes with a Ripple Neural Interface processor (Ripple Neuro; Salt Lake City, Utah) and sampled at 30,000 Hz, with 16 bits per sample and a user-selectable resolution of 0.25 or 0.50 µV per least-significant bit. We adjusted our grounding, amplifier, and stimulation settings to ensure that the recording system was not saturated and that signals remained within its amplifiers’ linear range (±12 mV at 0.5 µV resolution). The raw signal was initially bandpass filtered between 0.3 – 7,500 Hz by hardware filters in the recording hardware.

Signals were further processed offline to extract spiking activity. The recorded signals were always bandpass filtering between 700-5000 Hz. For TI-tACS and AM-tACS, notch filters were also applied at the carrier frequencies and their harmonics; for tACS, the stimulation artifact was already removed by the high-pass filtering. Next, spike thresholds were set at ±3σ, robustly estimated via the median absolute deviation. Snippets around each threshold crossing were extracted and clustered with UltraMegaSort2000, a k-means overclustering spike sorter(Hill et al., 2011). Its output was manually reviewed to ensure they had a consistent shape, a clear refractory period, and good separation in PCA space.

### Brain Stimulation

Electrical stimulation was delivered through two NeuroConn DC-STIMULATOR PLUS devices (NeuroConn, Ilmenau, Germany), modified by the manufacturer to produce high-frequency outputs, as described by Isak et al. (2023). The devices were operated in “remote mode”, where each converted a voltage input into a constant current output (1V = 2 mA peak-to-peak, or ±1 mA). The voltage inputs to both devices were provided by a dual channel arbitrary waveform (B&K Precision Model 4053B), which was programmed to emit sine waves, amplitude-modulated sine waves, or sweeps, as required. This, in turn, was triggered by TTL pulses from the stimulus control computer. Because the signal generator and stimulators have floating grounds, offsets between them can cause very small amounts of current to flow, even in the absence of a command signal. We therefore manually adjusted the devices to minimize this. We cannot formally exclude the possibility that very weak DC current was delivered in our experiments, but if so, it was less than ten percent of the weakest stimuli: 0.2 mA DC; ±1 mA (or 2 mA peak-to-peak) tACS.

Stimulation was applied to the scalp using 1 cm (radius) silver/silver chloride electrode (PISTIM; Neuroelectrics, Barcelona, Spain). Each electrode was coated with a conductive paste (SignaGel; Parker Labs Fairfield, New Jersey) and attached to the intact scalp with a biocompatible silicon elastomere (KwikSil; World Precision Instruments, Sarasota, FL]). Electrode impedances were typically 1-2 kOhms and never exceeded 15 kOhms.

To place the electrodes, we used finite-element modelling to identify suitable scalp locations for stimulating our target structures. In brief, each animal’s preoperative T1 and T2 MRIs were processed with SimNIBS 3.2.6 (Saturnino et al., 2021). Following automatic segmentation with the headreco pipeline and manual refinement, each element was classified as one of six tissue types: scalp, bone, CSF, eye, grey matter, or white matter and assigned the default conductivity used by SimNIBS. Simulated electrodes were attached to the model, using a 10-10 like grid to cover the scalp. Next, we calculated a leadfield by simulating the flow of current between a reference electrode at the vertex and each other scalp location. By combining appropriate elements of the leadfield, the electric fields resulting from stimulation of any two scalp locations can be calculated. A TI-tACS montage consists of two electrode pairs, each producing an electric field at a slightly different frequency. Using the leadfield, we exhaustively searched all possible configurations of four electrodes and calculated the modulation depth for each using the formula in Rampersad et al. (2019). From this list, we selected a montage predicted to produce a field of ∼0.7 V/m; the inset in Figure 1B shows predicted fields for one of our animals. However, this may be an overestimate, because the model relies on conductivities measured at lower frequencies, where current seems to penetrate more effectively (Figure 5). Since our electrodes are linear, and not aligned with the electric field, we could not measure the full three-dimensional vector field needed to calculate the overall field strength. However, these models have been extensively validated for conventional tACS (Huang et al., 2017; Jackson et al., 2016; Krause et al., 2019b).

To assess potential frequency-dependence, we also applied “chirp” or sweep stimuli through one pair of electrodes. In these experiments, the signal’s instantaneous frequency varied as a function of time, increasing from 10 to 2000 Hz over 30 s. We then calculated the instantaneous amplitude of stimulation artifact the chirp produced using a Hilbert transform, the results of which are shown in Figure 5.

For AM-tACS stimulation (Figure 6), we matched the modulation depth to that of the tACS stimulation used in the same experiments. Based on the data in Figure 5, we estimated that stimulation at 2000 Hz with ±2.5 mA of current would produce similar— and perhaps slightly stronger—modulation as the ±1.0 mA 10 Hz tACS. This was indeed the case in our experiments, where the AM-tACS modulation depth near our cells was between 1.14 and 1.17 times larger than the tACS. If anything, this should bias the comparison in favor of AM-tACS, but the results shown in Figure 6B nevertheless indicate that it was considerably weaker.

### Quantification and Statistical Analysis

Ethical and practical considerations limit the number of animals from which we can collect data. However, the critical comparisons in this paper are made within neurons (e.g., TI-tACS versus baseline or conventional tACS), making the cell rather than the animal the relevant unit of analysis (Fries & Maris, 2021). Where possible, statistical analyses used nonparametric tests to avoid distributional assumptions and all reported values are two-tailed. We did not carry out a formal power analysis because data characterizing TI-tACS effects in primates has not previously been reported. Sample sizes were instead determined based on our prior work with tACS, which allowed us to characterize changes in phase locking with 1.5 to 5 minutes of data. Data were analyzed using MATLAB (The MathWorks).

#### Neural activity

We quantified neural entrainment by calculating for each cell the pairwise phase consistency (PPC) value, a measure of the synchronization between the phase of a continuous signal (like the LFP or TI-tACS envelope) and a point process, single-unit spiking activity (Vinck et al., 2010). Because spikes can introduce small but detectable changes in signals recorded from the same electrode (Zanos et al., 2011), we compared spikes on one channel with the continuous signal from an adjacent channel 150 µm away. By using a local signal, rather a copy of the stimulator output, referencing conditions remain constant across conditions and any physiological distortion of the signal is incorporated into our measurements (Noury et al., 2016). During baseline, the continuous signal was the LFP, filtered in a ±1 Hz band around the stimulation frequency used in the rest of the experiment. During TI-tACS and AM-tACS, we first extracted the envelope of this signal (via Matlab’s envelope function) before filtering; this step was unnecessary for tACS. Thus, in all conditions, the signal reflects the electric potential in the extracellular space near each neuron. Next, we calculated the phase of the continuous signal via the Hilbert transform. Note that this correctly assigns a complete 360° of phase to each cycle of the AM. For each condition, we extracted the signal’s phase at the time of each spike and use these to calculate PPC values. These were compared across conditions via a randomization test. The PPC has several desirable statistical properties: notably, it is unbiased and unaffected by the number of spikes. However, PPC values are rarely reported in the literature, so we transformed them to phase locking values (PLV) by taking their square root. The resulting PLVs are also equivalent (under some simplifying assumptions) to spike-field coherence, another oft-used metric of spike-LFP coupling. Linear mixed-effects models, with a fixed effect of stimulation and random effect of block, were used to assess firing rates for individual neurons.

#### Factors Affecting TI-tACS Response

The data in Figure 3 suggest that baseline levels of entrainment predict the subsequent effects of TI-tACS. If the neurons’ PLVs were fluctuating randomly, regression to the mean could cause a similar pattern of effects: cells which had a spuriously high PLV in one condition would have a lower one in the other (and vice versa). However, there are several theoretical and statistical reasons to believe this is not occurring in our data.

First, this outcome is consistent with prior work using conventional tACS as well as the mathematical properties of interacting oscillators (Krause et al. 2022). Second, we find statistically significant effects at both the single-cell and population levels. Although regression to the mean could conceivably produce the population-level result, random fluctuations in the activity of individual cells are—by definition—unlikely to be statistically significant. In our sample, 28% of the neurons (65/234 cells) exhibited statistically significant changes, which is extremely unlikely assuming at our per-cell threshold of 0.05 (p<0.0001; Fisher’s exact test). Inverse correlations between the baseline and TI-tACS PLVs remained significant even after applying Oldham’s Method (Oldham, 1962) to correct for regression to the mean (Spearman’s ⍴=-0.43; p << 0.001). We also analyzed the data with ANCOVAs, which protect against regression the mean (Barnett et al., 2005; Clifton & Clifton, 2019). In these models, baseline entrainment, amplitude, and their interactions were the only significant predictors of either the change score (i.e., ΔPLV) or the TI-tACS PLV. The other two factors (brain area and AM frequency) were not statistically significant during the ANOCOVAs. A model comparison approach also found that these factors were not parsimonious components of the model, as they led to only minor changes in ΔAIC. For firing rates, a similar analysis found that only the baseline firing rate predicted activity during TI-tACS blocks.

### Technical Valida1on

We performed a series of control experiments and analyses to exclude potentially artifactual results.

First, prior work has found that the high frequency carriers used during TI-tACS and AM-tACS can produce artifacts at the modulation frequency (i.e., 20 Hz during 2000 + 2020 Hz TI-tACS), due to non-linear transfer characteristics in either the stimulation or recording system (Kasten et al., 2018). We therefore tested both systems with a saline bath phantom. This experiment included the complete signal pathway, beginning with the same electrodes used in our experiments and ending with the same digitization and stimulation systems. No low-frequency artifacts were detectable in our apparatus (Figure S1). Moreover, our readout is the timing of action potentials, which have a characteristic shape that would be hard to confuse with the sinusoidal stimulation artifacts.

However, the carrier frequencies do overlap spectrally with spike frequency band. A loss of single-unit isolation could potentially produce apparent changes in entrainment. However, there are several reasons to believe this does not happen in our data. First, the overall firing rate varied little between conditions (∼0.1 Hz; see Results), so a very particular pattern of artifactual insertions and deletions would be required to preserve the rate while simulating changes in spike timing. These artifacts would also have different shapes from true spike waveforms, but in our data, the shape of the spikes remained consistent across conditions (minimum r=0.95; see Figures 2 and 4 for examples). We also validated our spike sorting pipeline by adding 2000 Hz noise to a segment of baseline data and resorting it. Outcomes were similar between conditions (1509 spikes detected in both conditions, correlation between waveforms: r=0.999). In short, we consistently detected spikes even when high-frequency stimulation artifacts were present, presumably because spikes, which are relatively sharp, have a relatively broad spectrum and so their shape is only minimally affected by very narrowband artifact removal.

Attenuation of high frequencies by our recording apparatus could also produce the apparent fall-off seen in Figure 5. Our headstages do contain hardware anti-aliasing filters: a 3rd-order Butterworth filter with a 7500 Hz corner frequency. However, theoretical calculations indicate that such a filter should produce ∼0.08 dB (or 2%) attenuation at 2000 Hz. We confirmed this calculation by directly connecting the signal generator outputs to the headstage, which also demonstrated very little attenuation during frequency sweeps, suggesting a biological origin (Figure 5, grey line).

In a similar vein, TI-tACS could appear weak if current were shunted between the two stimulators, rather than entering the brain. However, this explanation is hard to square with the results of the AM-tACS experiments, where only a single pair of electrodes were used and the possibility of shunting was therefore physically eliminated.

## Supplemental Figures

**Figure S1.**
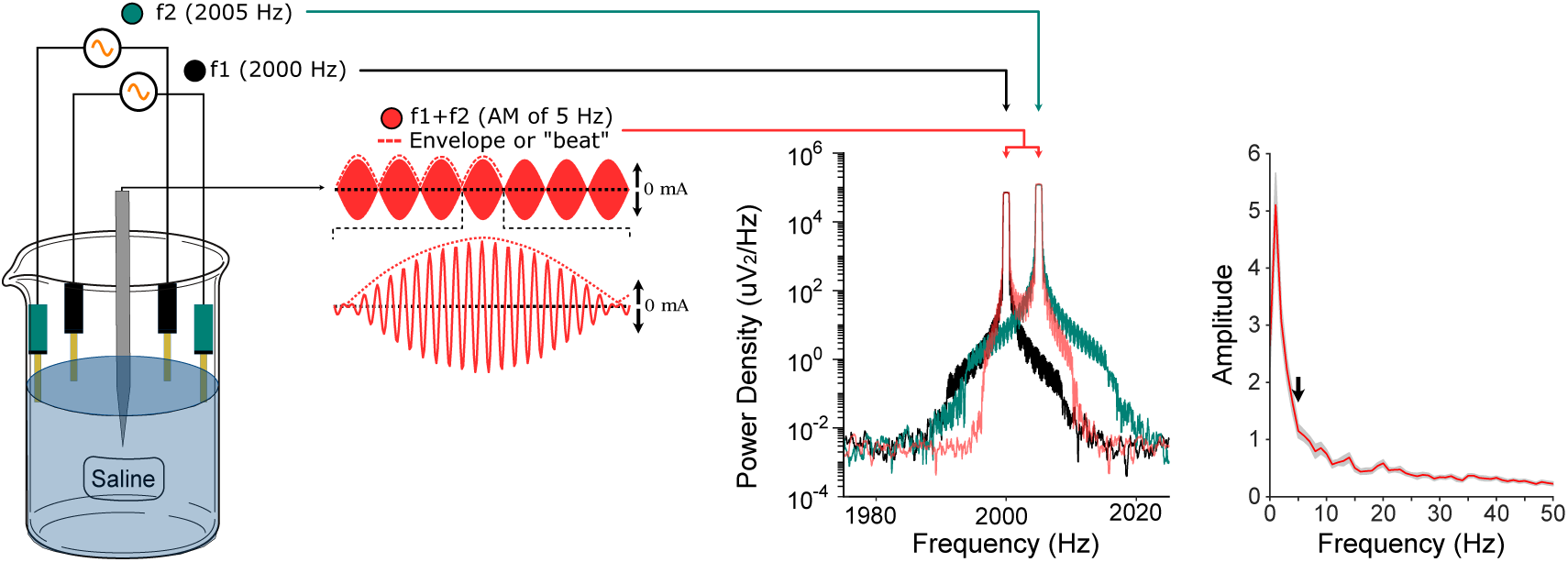
Validation experiment. Recordings from a saline bath confirm that no low-frequency artifacts, especially at the AM frequency (red, inset arrow), were produced by our stimulation and recording systems. Only components at the two carrier frequencies (green and black) were present in these data.

**Figure S2.**
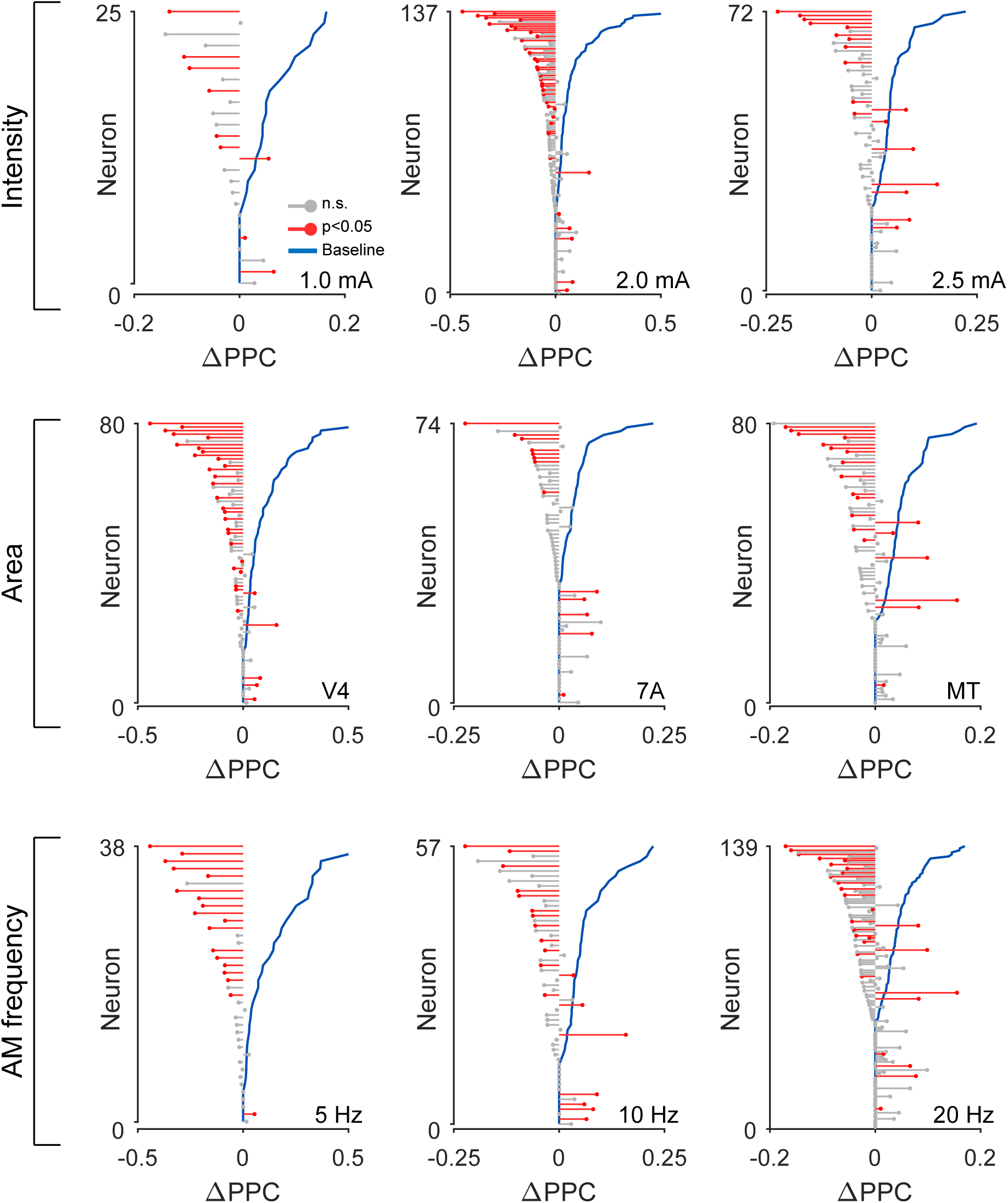
Data from Main Figure 3, plotted in the same style, but separated by stimulation amplitude (top row), brain area (middle), or AM frequency (bottom). As described in the main text, these parameters were not significant predictors of ΔPLV. Instead, the baseline PLV (blue) determines the effects of subsequent stimulation ΔPLV.

## References

1. Asamoah, B., Khatoun, A., & Laughlin, M. M. (2022). Frequency-Specific Modulation of Slow-Wave Neural Oscillations via Weak Exogeneous Extracellular Fields Reveals a Resonance Pattern. Journal of Neuroscience, 42(32), 6221–6231. 10.1523/JNEUROSCI.0177-22.2022

2. Baker, C. L., & Mareschal, I. (2001). Chapter 12 Processing of second-order stimuli in the visual cortex. In Progress in Brain Research (Vol. 134, pp. 171–191). Elsevier. 10.1016/S0079-6123(01)34013-X

3. Barnett, A. G., van der Pols, J. C., & Dobson, A. J. (2005). Regression to the mean: What it is and how to deal with it. International Journal of Epidemiology, 34(1), 215–220. 10.1093/ije/dyh299

4. Budde, R. B., Williams, M. T., & Irazoqui, P. P. (2023). Temporal interference current stimulation in peripheral nerves is not driven by envelope extraction. Journal of Neural Engineering, 20(2), 026041. 10.1088/1741-2552/acc6f1

5. Cassarà, A. M., Newton, T. H., Zhuang, K., Regel, S. J., Achermann, P., Kuster, N., & Neufeld, E. (2022). Safety Recommendations for Temporal Interference Stimulation in the Brain (p. 2022.12.15.520077). bioRxiv. 10.1101/2022.12.15.520077

6. Chaieb, L., Antal, A., & Paulus, W. (2011). Transcranial alternating current stimulation in the low kHz range increases motor cortex excitability. Restorative Neurology and Neuroscience, 29(3), 167–175. 10.3233/RNN-2011-0589

7. Clifton, L., & Clifton, D. A. (2019). The correlation between baseline score and post-intervention score, and its implications for statistical analysis. Trials, 20(1), 43. 10.1186/s13063-018-3108-3

8. Cohen, M. R., & Maunsell, J. H. R. (2009). Attention improves performance primarily by reducing interneuronal correlations. Nature Neuroscience, 12(12), 1594–1600. 10.1038/nn.2439

9. Esmaeilpour, Z., Kronberg, G., Reato, D., Parra, L. C., & Bikson, M. (2021). Temporal interference stimulation targets deep brain regions by modulating neural oscillations. Brain Stimulation, 14(1), 55–65. 10.1016/j.brs.2020.11.007

10. Fries, P., & Maris, E. (2021). What to do if N is two? (arXiv:2106.14562). arXiv. 10.48550/arXiv.2106.14562

11. Grossman, N., Bono, D., Dedic, N., Kodandaramaiah, S. B., Rudenko, A., Suk, H.-J., Cassara, A. M., Neufeld, E., Kuster, N., Tsai, L.-H., Pascual-Leone, A., & Boyden, E. S. (2017). Noninvasive Deep Brain Stimulation via Temporally Interfering Electric Fields. Cell, 169(6), 1029–1041.e16. 10.1016/j.cell.2017.05.024

12. Hasgall, P., Neufeld, E., Gosselin, M.-C., Klingenböck, A., & Kuster, N. (2022). ITIS Database for thermal and electromagnetic parameters of biological tissues, Version 2.2. 4.1. https://itis.swiss/virtual-population/tissue-properties/downloads/database-v4-1/

13. Hill, D. N., Mehta, S. B., & Kleinfeld, D. (2011). Quality metrics to accompany spike sorting of extracellular signals. The Journal of Neuroscience, 31(24), 8699–8705. 10.1523/JNEUROSCI.0971-11.2011

14. Huang, Y., Liu, A. A., Lafon, B., Friedman, D., Dayan, M., Wang, X., Bikson, M., Doyle, W. K., Devinsky, O., & Parra, L. C. (2017). Measurements and models of electric fields in the in vivo human brain during transcranial electric stimulation. ELife, 6, e18834. 10.7554/eLife.18834

15. Huang, Y., & Parra, L. C. (2019). Can transcranial electric stimulation with multiple electrodes reach deep targets? Brain Stimulation, 12(1), 30–40. 10.1016/j.brs.2018.09.010

16. Iszak, K., Gronemann, S. M., Meyer, S., Hunold, A., Zschüntzsch, J., Bähr, M., Paulus, W., & Antal, A. (2023). Why Temporal Inference Stimulation May Fail in the Human Brain: A Pilot Research Study. Biomedicines, 11(7), Article 7. 10.3390/biomedicines11071813

17. Jackson, M. P., Rahman, A., Lafon, B., Kronberg, G., Ling, D., Parra, L. C., & Bikson, M. (2016). Animal models of transcranial direct current stimulation: Methods and mechanisms. Clinical Neurophysiology, 127(11), 3425–3454. 10.1016/j.clinph.2016.08.016

18. Johnson, L., Alekseichuk, I., Krieg, J., Doyle, A., Yu, Y., Vitek, J., Johnson, M., & Opitz, A. (2020). Dose-dependent effects of transcranial alternating current stimulation on spike timing in awake nonhuman primates. Science Advances, 6(36), eaaz2747. 10.1126/sciadv.aaz2747

19. Kasten, F. H., Negahbani, E., Fröhlich, F., & Herrmann, C. S. (2018). Non-linear transfer characteristics of stimulation and recording hardware account for spurious low-frequency artifacts during amplitude modulated transcranial alternating current stimulation (AM-tACS). NeuroImage, 179, 134–143. 10.1016/j.neuroimage.2018.05.068

20. Khalifa, A., Abrishami, S. M., Zaeimbashi, M., Tang, A. D., Coughlin, B., Rodger, J., Sun, N. X., & Cash, S. S. (2023). Magnetic temporal interference for noninvasive and focal brain stimulation. Journal of Neural Engineering, 20(1), 016002. 10.1088/1741-2552/acb015

21. Khatoun, A., Asamoah, B., & Mc Laughlin, M. (2019). How does transcranial alternating current stimulation entrain single-neuron activity in the primate brain? Proceedings of the National Academy of Sciences, 116(45), 22438–22439. 10.1073/pnas.1912927116

22. Kilgore, K. L., & Bhadra, N. (2014). Reversible Nerve Conduction Block Using Kilohertz Frequency Alternating Current. Neuromodulation: Journal of the International Neuromodulation Society, 17(3), 242–255. 10.1111/ner.12100

23. Krause, M. R., Vieira, P. G., Csorba, B. A., Pilly, P. K., & Pack, C. C. (2019a). Reply to Khatoun et al.: Speculation about brain stimulation must be constrained by observation. Proceedings of the National Academy of Sciences, 116(45), 22440– 22441. 10.1073/pnas.1914483116

24. Krause, M. R., Vieira, P. G., Csorba, B. A., Pilly, P. K., & Pack, C. C. (2019b). Transcranial alternating current stimulation entrains single-neuron activity in the primate brain. Proceedings of the National Academy of Sciences, 116(12), 5747– 5755. 10.1073/pnas.1815958116

25. Krause, M. R., Vieira, P. G., Thivierge, J.-P., & Pack, C. C. (2022). Brain stimulation competes with ongoing oscillations for control of spike timing in the primate brain. PLOS Biology, 20(5), e3001650. 10.1371/journal.pbio.3001650

26. Kuzin, M., Sachkov, V., & Zhukovsky, V. (1965). Electronarcosis produced by interference currents in clinical practice. Present. Tech. Surg. Data Sixth Sci. Sess. Sci. Res. Inst. Exp. Surg. Appar. Instruments JPRS, 31, 4–5.

27. Labruna, L., Merrick, C., Peterchev, A. V., Inglis, B., Ivry, R. B., & Sheltraw, D. (2023). Kilohertz Transcranial Magnetic Perturbation (kTMP): A New Non-invasive Method to Modulate Cortical Excitability (p. 2021.11.17.465477). bioRxiv. 10.1101/2021.11.17.465477

28. Liu, A., Vöröslakos, M., Kronberg, G., Henin, S., Krause, M. R., Huang, Y., Opitz, A., Mehta, A., Pack, C. C., Krekelberg, B., Berényi, A., Parra, L. C., Melloni, L., Devinsky, O., & Buzsáki, G. (2018). Immediate neurophysiological effects of transcranial electrical stimulation. Nature Communications, 9(1), 5092. 10.1038/s41467-018-07233-7

29. Luff, C. E., Peach, R., Mallas, E.-J., Rhodes, E., Laumann, F., Boyden, E. S., Sharp, D. J., Barahona, M., & Grossman, N. (2023). The neuron mixer and its impact on human brain dynamics (p. 2023.01.05.522833). bioRxiv. 10.1101/2023.01.05.522833

30. McConnell, G. C., So, R. Q., Hilliard, J. D., Lopomo, P., & Grill, W. M. (2012). Effective deep brain stimulation suppresses low-frequency network oscillations in the basal ganglia by regularizing neural firing patterns. The Journal of Neuroscience, 32(45), 15657–15668. 10.1523/JNEUROSCI.2824-12.2012

31. Metzen, M. G., & Chacron, M. J. (2015). Neural Heterogeneities Determine Response Characteristics to Second-, but Not First-Order Stimulus Features. Journal of Neuroscience, 35(7), 3124–3138. 10.1523/JNEUROSCI.3946-14.2015

32. Mirzakhalili, E., Barra, B., Capogrosso, M., & Lempka, S. F. (2020). Biophysics of Temporal Interference Stimulation. Cell Systems, 11(6), 557–572.e5. 10.1016/j.cels.2020.10.004

33. Noury, N., Hipp, J. F., & Siegel, M. (2016). Physiological processes non-linearly affect electrophysiological recordings during transcranial electric stimulation. NeuroImage, 140, 99–109. 10.1016/j.neuroimage.2016.03.065

34. Oldham, P. D. (1962). A note on the analysis of repeated measurements of the same subjects. Journal of Chronic Diseases, 15(10), 969–977. 10.1016/0021-9681(62)90116-9

35. Perlmutter, J. S., & Mink, J. W. (2006). Deep brain stimulation. Annual Review of Neuroscience, 29, 229–257. 10.1146/annurev.neuro.29.051605.112824

36. Plovie, T., Schoeters, R., Tarnaud, T., Martens, L., Joseph, W., & Tanghe, E. (2023). Nonlinearities and Timescales in Temporal Interference Stimulation (p. 2022.02.04.479138). bioRxiv. 10.1101/2022.02.04.479138

37. Radman, T., Ramos, R. L., Brumberg, J. C., & Bikson, M. (2009). Role of cortical cell type and morphology in subthreshold and suprathreshold uniform electric field stimulation in vitro. Brain Stimulation, 2(4), 215–228, 228.e1-3. 10.1016/j.brs.2009.03.007

38. Radman, T., Su, Y., An, J. H., Parra, L. C., & Bikson, M. (2007). Spike timing amplifies the effect of electric fields on neurons: Implications for endogenous field effects. The Journal of Neuroscience, 27(11), 3030–3036. 10.1523/JNEUROSCI.0095-07.2007

39. Rampersad, S., Roig-Solvas, B., Yarossi, M., Kulkarni, P. P., Santarnecchi, E., Dorval, A. D., & Brooks, D. H. (2019). Prospects for transcranial temporal interference stimulation in humans: A computational study. NeuroImage, 202, 116124. 10.1016/j.neuroimage.2019.116124

40. Rosenberg, A., & Issa, N. P. (2011). The Y cell visual pathway implements a demodulating nonlinearity. Neuron, 71(2), 348–361. 10.1016/j.neuron.2011.05.044

41. Saturnino, G. B., Madsen, K. H., & Thielscher, A. (2021). Optimizing the electric field strength in multiple targets for multichannel transcranial electric stimulation. Journal of Neural Engineering, 18(1), 014001. 10.1088/1741-2552/abca15

42. Seidlitz, J., Sponheim, C., Glen, D., Ye, F. Q., Saleem, K. S., Leopold, D. A., Ungerleider, L., & Messinger, A. (2018). A population MRI brain template and analysis tools for the macaque. NeuroImage, 170, 121–131. 10.1016/j.neuroimage.2017.04.063

43. Shofner, W. P., Sheft, S., & Guzman, S. J. (1996). Responses of ventral cochlear nucleus units in the chinchilla to amplitude modulation by low-frequency, two-tone complexes. The Journal of the Acoustical Society of America, 99(6), 3592– 3605. 10.1121/1.414957

44. Thiele, C., Zaehle, T., Haghikia, A., & Ruhnau, P. (2021). Amplitude modulated transcranial alternating current stimulation (AM-TACS) efficacy evaluation via phosphene induction. Scientific Reports, 11(1), 22245. 10.1038/s41598-021-01482-1

45. Venables, N. C., Bernat, E. M., & Sponheim, S. R. (2009). Genetic and Disorder-Specific Aspects of Resting State EEG Abnormalities in Schizophrenia. Schizophrenia Bulletin, 35(4), 826–839. 10.1093/schbul/sbn021

46. Vieira, P. G., Krause, M. R., & Pack, C. C. (2020). TACS entrains neural activity while somatosensory input is blocked. PLOS Biology, 18(10), e3000834. 10.1371/journal.pbio.3000834

47. Vinck, M., van Wingerden, M., Womelsdorf, T., Fries, P., & Pennartz, C. M. A. (2010). The pairwise phase consistency: A bias-free measure of rhythmic neuronal synchronization. NeuroImage, 51(1), 112–122. 10.1016/j.neuroimage.2010.01.073

48. Violante, I. R., Alania, K., Cassarà, A. M., Neufeld, E., Acerbo, E., Carron, R., Williamson, A., Kurtin, D. L., Rhodes, E., Hampshire, A., Kuster, N., Boyden, E. S., Pascual-Leone, A., & Grossman, N. (2022). Non-invasive temporal interference electrical stimulation of the human hippocampus (p. 2022.09.14.507625). bioRxiv. 10.1101/2022.09.14.507625

49. Vöröslakos, M., Takeuchi, Y., Brinyiczki, K., Zombori, T., Oliva, A., Fernández-Ruiz, A., Kozák, G., Kincses, Z. T., Iványi, B., Buzsáki, G., & Berényi, A. (2018). Direct effects of transcranial electric stimulation on brain circuits in rats and humans. Nature Communications, 9(1), Article 1. 10.1038/s41467-018-02928-3

50. Zanos, T. P., Mineault, P. J., & Pack, C. C. (2011). Removal of Spurious Correlations Between Spikes and Local Field Potentials. Journal of Neurophysiology, 105(1), 474–486. 10.1152/jn.00642.2010

